# Targeting MYC Overexpressing Leukemia with Cardiac Glycoside Proscillaridin Through Downregulation of Histone Acetyltransferases

**DOI:** 10.1101/444133

**Authors:** Elodie M. Da Costa, Gregory Armaos, Gabrielle McInnes, Annie Beaudry, Gaël Moquin-Beaudry, Virginie Bertrand-Lehouillier, Maxime Caron, Chantal Richer, Pascal St-Onge, Jeffrey R. Johnson, Nevan Krogan, Yuka Sai, Michael Downey, Moutih Rafei, Meaghan Boileau, Kolja Eppert, Ema Flores-Díaz, André Haman, Trang Hoang, Daniel Sinnett, Christian Beauséjour, Serge McGraw, Noël J-M Raynal

## Abstract

Targeting MYC oncogene remains a major therapeutic goal in cancer chemotherapy. Here, we demonstrate that proscillaridin, a cardiac glycoside approved for heart failure treatment exhibit anticancer selectivity towards high MYC expressing leukemic cell lines and leukemia stem cells. At a clinically relevant concentration, proscillaridin induced a rapid downregulation of MYC protein level, due to a significant decrease in MYC protein half-life. Proscillaridin treatment induced a downregulation of gene sets involved in MYC pathway, and a concomitant upregulation of genes involved in hematopoietic differentiation. Proscillaridin induced a significant loss of lysine acetylation in histone H3 (K9, K14, K18 and K27) and in non-histone proteins such as MYC, MYC target proteins, and a series of histone acetylation regulators. Loss of lysine acetylation correlated with a rapid downregulation of histone acetyltransferase protein levels, involved in histone and MYC acetylation (CBP, P300, GCN5, TIP60, and MOZ), preferentially in MYC overexpressing leukemia as compared to other cancer cells. These results support the repurposing of proscillaridin in MYC overexpressing leukemia and propose a novel strategy to target MYC in cancer.

## INTRODUCTION

MYC (c-MYC) transcription factor is a major driver of oncogenic transcriptional programs. It contributes to gene deregulation in cancer by promoting expression of genes involved in cell proliferation (1). High MYC expression drives tumor initiation, progression, and maintenance and is associated with aggressive cancers and poor prognoses (2,3). MYC is a well-known driver in leukemia inducing cell proliferation and blocking cell differentiation (4). Moreover, MYC contributes to long-term self-renewal of leukemic stem cells (5). Genetic suppression of MYC in transgenic mouse models induces differentiation and cell growth arrest of leukemic cells (6-8). Therefore, targeting MYC addiction in leukemia is a major therapeutic goal. Since MYC lacks a catalytic site, its direct inhibition has been extremely challenging. Indirect MYC inhibition demonstrated therapeutic efficacy with bromodomain inhibitors (such as JQ1 or THZ1), by blocking MYC transcriptional effects (9-13). Unfortunately, cancer cells, such as leukemia, breast and ovarian cancers, develop resistance to these inhibitors by compensatory mechanisms using other bromodomain containing proteins or kinome reprogramming (14-16). Together, these studies highlight the need to develop new strategies to abrogate MYC addiction in cancer.

MYC protein stability is regulated by post-translational modifications and acetylation increases its half-life (17,18). The deposition of acetyl groups on MYC lysine residues is catalyzed by lysine acetyltransferases (KATs) such as CBP (KAT3A), P300 (KAT3B), GCN5 (KAT2A) and Tip60 (KAT5) KATs acetylate also histone proteins causing chromatin opening and gene activation (19). Lysine acetyltransferase pharmacological inhibition represent an interesting strategy to target indirectly MYC by blocking upstream mechanisms involved in its stability. However, KATs have overlapping targets and commercially available KAT inhibitors require further optimization (20).

While screening more than 1,000 FDA-approved drugs, we reported that cardiac glycosides, which are approved for heart failure treatment, exhibit significant epigenetic and anticancer effects (21,22). Cardiac glycosides, including digitoxin, digoxin, lanatoside, ouabain and proscillaridin, triggered reactivation of epigenetically silenced tumor suppressor genes (22). Moreover, all cardiac glycosides produced synergistic responses when used in combination with the epigenetic drug decitabine (demethylating agent) (21). Several epidemiological studies argue in favor of repurposing cardiac glycosides in oncology. Notably, patients treated with cardiac glycosides for heart failure present a lower rate of cancer diagnosis as compared to the general population (23). Upon cancer diagnosis, these patients have generally a less aggressive disease and respond better to therapy (23,24).

Repurposing cardiac glycosides in oncology is limited by their narrow therapeutic window, for which maximal plasmatic level is around 10 nanomolar (nM), causing cardiac toxicities due to Na^+^/K^+^ pump inhibition (23,25-27). Several *in vitro* and *in vivo* studies tested their anticancer activity at supra-pharmacological doses, which are not reachable in humans; in particular, in rodents who can tolerate high doses of these drugs due to structural differences in Na^+^/K^+^ pump as compared to human (26,28,29). Since the repurposing of cardiac glycosides is restricted to the low nanomolar range, we sought to identify cancer types highly sensitive to these drugs. To do so, we screened a panel of 14 human cancer cell lines with proscillaridin, which was identified as the most potent cardiac glycoside in our previous screens (21,22). Proscillaridin produced antiproliferative effects with a preferential selectivity towards MYC overexpressing leukemia cells. We demonstrated that proscillaridin produced a global loss of acetylation in chromatin and MYC itself, producing epigenetic effects and MYC downregulation. These results provide compelling evidence for the repurposing of cardiac glycoside proscillaridin against leukemia driven by MYC oncogenic signature.

## MATERIALS AND METHODS

### Cell Culture and Drug Treatments

A panel of 14 human cancer cell lines and *hTERT/SV40ER-*immortalized human primary fibroblasts transformed with *MYC, RAS*^*V12*^ or *MYC* and *RAS*^*V12*^ were used in the study. Cell types and culture conditions are described in Supplementary Materials and Methods. Proscillaridin was purchased from Santa Cruz Biotechnologies. IC_50_ values were calculated with GraphPad Prism software. For cycloheximide-chase assay, MOLT-4 cells were treated with cycloheximide (150 µg/ml) purchased from Acros Organics.

### Protein and Histone Extractions

Whole cell proteins were extracted using cold whole-cell lysis buffer (50 mM Tris-Cl pH 7.4, 5 mM EDTA, 250 mM NaCl, 50 mM NaF, 0.1% Triton, 0.1 mM Na_3_VO_4_, and 1 mM PMSF), supplemented with Complete™ Protease Inhibitor Cocktail (Roche). Histones were harvested using acid-extraction method with cold Triton Extraction Buffer (TEB; 0.5% Triton, 2 mM PMSF, 0.02% NaN_3_, 10 mM sodium butyrate), supplemented with protease inhibitor cocktail. Protein extracts were separated by SDS-PAGE and transferred onto a polyvinyl difluoride membrane. All experiments were performed in triplicate. Antibodies are listed in the Supplemental Materials and Methods section.

### RNA Extraction, Sequencing and Analysis

QIAshredder was used to homogenize cell lysates and eliminate debris prior to RNA extraction using RNeasy Mini Kit. Briefly, 10 µg of purified RNA was treated with DNAse and quantified by Agilent RNA 6,000 Nano kit bioanalyser chips. 1 µg of mRNA was used for library preparation with TruSEq Stranded mRNA LT. RNA sequencing was performed using HiSeq 2500. Experiments were performed in triplicate. Reads were aligned to human genome (hg19) using STAR v2.4.2 and differential gene expression analysis between untreated and treated cells was done using DESeq2 v1.10.1 (30,31). For bioinformatics analyses, data were processed using gene set enrichment analyses (GSEA, broadinstitute.org/gsea), metascape (metascape.org) and gene mania (genemania.org). MOLT-4 cells H3K27ac ChIP-seq data from publicly available dataset (GEO: GSM2037790) were used in associated with transcriptomic data. GSEA analysis of 8227 AML fractions and the LSC signatures was performed using the control sample data from GSE55814. GEO2R was used to generate a ranked list of LSC-related genes (6 LSC CD34^+^CD38^-^ samples vs 12 non-LSC CD34^-^ samples) used in GSEA analysis.

### Acetylation Analysis by Immunoprecipitation and Mass Spectrometry

Whole cell protein extracts were incubated overnight with 5 µg/ml of MYC antibody (Abcam, AB32072). After immunoprecipitation and transfer, proteins were probed with lysine pan-acetyl antibody (1:2500 Cell Signaling, 9681). For acetylome analysis by mass spectrometry, samples were prepared as previously described (32). Briefly, 4 biological replicates of untreated and proscillaridin-treated MOLT-4 cells (5 nM, 48h) were digested with trypsin. Peptides were analyzed by mass-spectrometry and data were extracted with the MaxQuant software package (version 1.5.5.1) and subsequently analyzed using an in-house computational pipeline for statistical analysis of relative quantification with fixed and/or mixed effect models, implemented in the MSstats Bioconductor package (version 3.3.10) (33,34). Peptides were searched with SwissProt human protein database.

## RESULTS

### Cardiac Glycoside Proscillaridin Targets MYC-Driven Leukemic Cells

To identify cancer types with high sensitivity to proscillaridin at concentrations within its therapeutic window, we screened a panel of 14 human cancer cell lines and measured cancer cell proliferation after a 24h treatment. Upon calculating the half-maximal inhibitory concentration (IC_50_), we noticed a 2,800-fold difference in IC_50,_ with leukemic cell lines being more sensitive to proscillaridin (Supplementary Figure 1a). We hypothesized that this striking difference was due to the oncogene landscape driving each cancer cell type, which may influence drug activity. Since proscillaridin demonstrated greater activity against leukemia cell lines, we investigated whether expression of MYC protein, a major oncogenic driver in leukemia, would be associated with drug efficacy. There was a significant inverse correlation between MYC protein expression in untreated cancer cells and proscillaridin IC_50_ values (p = 0.0172; Supplementary Figure 1b). We identified MOLT-4 (acute lymphoblastic T-cell leukemia) and NALM-6 (acute lymphoblastic B-cell leukemia) cells as the most sensitive cell lines expressing high levels of MYC proteins, and two resistant cell lines (SW48-colorectal and A549-lung cancers) with low MYC levels (Figure 1a).

**Fig. 1.**
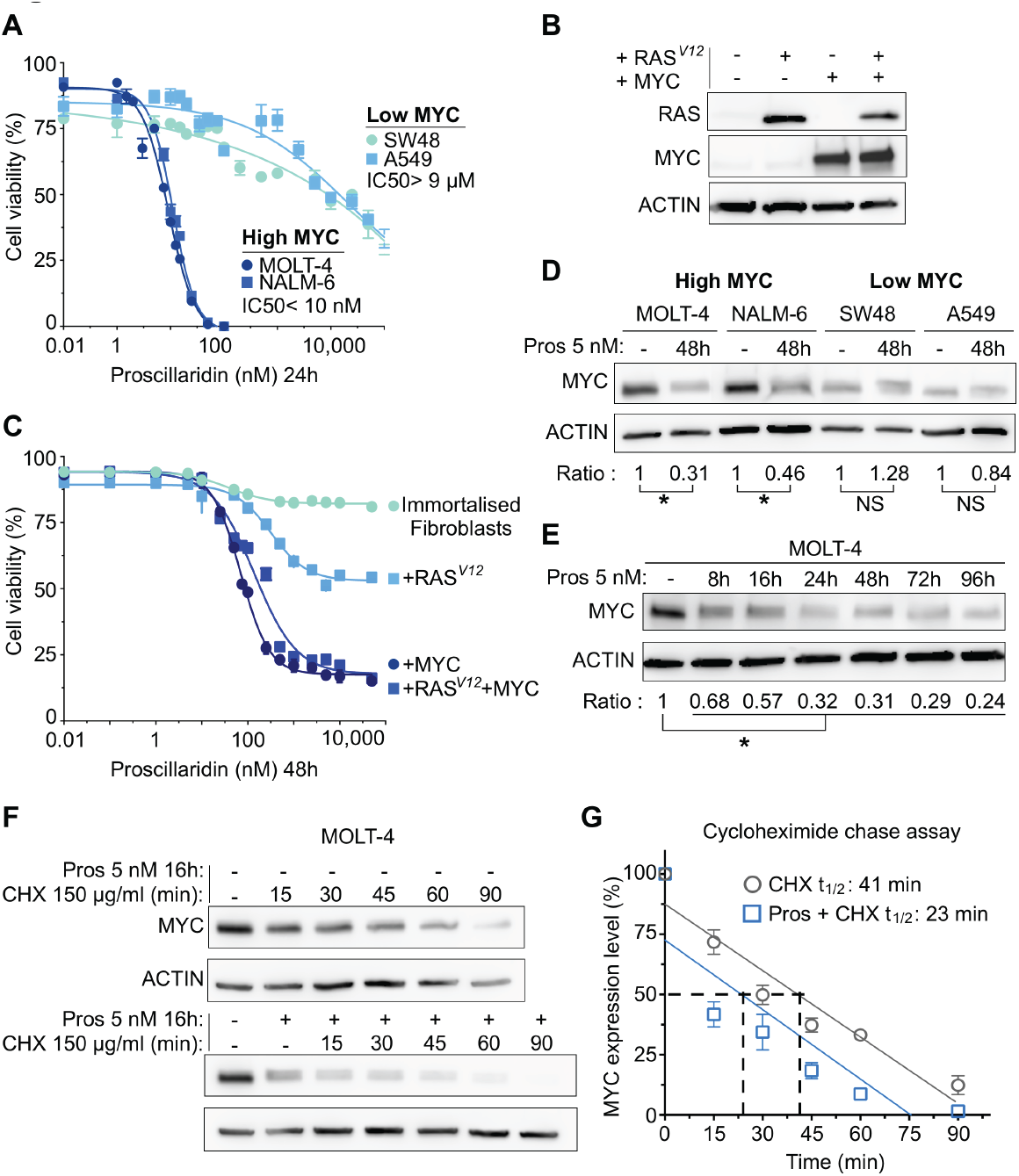
Targeting High MYC Expressing Cancer Cells with Cardiac Glycoside Proscillaridin. **a** Cell viability and half-maximal inhibitory concentration (IC_50_) calculations after a 24h proscillaridin treatment (ranging from 1 nM to 100 µM) in a high MYC expressing human leukemia cell lines (MOLT-4 and NALM-6) and in low MYC expressing human cancer cell lines (SW48 and A549) (n=4). **b** *MYC* and *RAS* protein expression assessed by western blotting in immortalized fibroblasts and fibroblasts transfected with *RAS*^*V12*^, *MYC* and *RAS*^*V12*^/*MYC* (n=3). **c** Dose response curves after 48h proscillaridin treatment (0.01 nM to 50 µM) in immortalized fibroblasts and fibroblasts transfected with *RAS*^*V12*^, *MYC* and *RAS*^*V12*^/*MYC* (n = 4). **d** MYC protein expression after proscillaridin treatment (5 nM; 48h) in MOLT-4, NALM-6, SW48 (colon cancer) and A549 (lung cancer) cells assessed by western blotting. MYC expression is calculated as a ratio over ACTIN levels (*indicates P<0.05; ANOVA; n = 3). **e** Time course experiment in MOLT-4 cells treated with proscillaridin at 5 nM (8h to 96h). MYC protein expression is calculated as a ratio over ACTIN levels (*indicates P<0.05; ANOVA; n = 3). **f** Effect of proscillaridin on MYC half-life in MOLT-4 cells. MYC half-life was estimated by cycloheximide (CHX, 150 µg/ml) chase assay treatment in MOLT-4 cells pretreated or not with proscillaridin (5 nM, 16h). MYC protein expression was performed by Western blotting and ACTIN expression was used as loading control (shown is representative blots of 3 independent experiments). **g** MYC expression level after cycloheximide treatment was quantified over ACTIN levels. Expression was calculated relative to the level of untreated cells (grey) or relative to the level following proscillaridin treatment (5 nM, 16h; blue) in MOLT-4 cells. Linear regression analysis was conducted and MYC half-life was calculated (n = 3).

To evaluate MYC’s contribution in cancer cell sensitivity to proscillaridin, we investigated drug response using an isogenic cell system consisting of *hTERT/SV40ER-*immortalized human primary fibroblasts, transformed with different oncogenes including *MYC, RAS*^*V12*^ or the combination of both oncogenes. This system allowed exploring the effect of oncogenic transformation within the same genetic background. We choose this approach as opposed to MYC overexpression or knockout genetic manipulations in leukemic cell lines since MYC overexpression amplifies more robustly MYC-dependent genes, and MYC knockout in leukemia leads to proliferation arrest, and cell death (6,35,36). After transfection, *MYC*-transformed fibroblasts had a small and round phenotype whereas *RAS*^*V12*^*-* transformed cells displayed increased vacuole formation and large cytoplasm. *MYC* and *RAS*^*V12*^-transformed cells exhibited a round phenotype with vacuole formation in the cytoplasm (Supplementary Figure 1c). High levels of MYC and RAS protein levels were detected after transfections when compared to non-transfected cells (Figure 1b). Using a wide range of proscillaridin concentrations (from 0.01 nM to 100 µM) for 48h, we measured cell viability and calculated IC_50_ values (Figure 1c). Untransformed fibroblasts were fully resistant to proscillaridin. Likewise, *RAS*^*V12*^ transformed cells were mildly affected by the treatment where proscillaridin at high doses failed to impact cell viability by more than 50%. Conversely, *MYC* transformed fibroblasts were highly sensitive to proscillaridin with an IC_50_ value of 70 nM. Moreover, *MYC* and *RAS*^*V12*^ transformed fibroblasts (referred as to *RAS*^*V12*^*+MYC*) had a low IC_50_ value (132 nM) despite the presence of *RAS*^*V12*^. Consequently, *MYC* overexpression was driving proscillaridin sensitivity in transformed fibroblasts. In comparison, after 48h treatment, MOLT-4 and NALM-6 cells (MYC overexpressing leukemic cells) showed IC_50_ values of 2.3 nM and 3 nM, respectively, which correspond to clinically achievable concentrations (Supplementary Figure 1d).

To explore the mechanism by which *MYC* overexpression in cancer cells correlates with proscillaridin sensitivity, we compared its effects between *MYC* driven leukemic cells (MOLT-4 and NALM-6) and low expressing *MYC* cancers driven by *KRAS* mutations (SW48 colon and A549 lung cancer cells). We found that proscillaridin (5 nM; 48h) significantly reduced MYC protein level by more than 50% in MOLT-4 and NALM-6 cells but not in SW48 and A549 cells (Figure 1d). Time-course experiments with both leukemic cell lines (8h to 96h) showed that proscillaridin induced a significant (up to 80%) and rapid MYC downregulation (Figure 1d; Supplementary Figure 1e). Lastly, we demonstrated that proscillaridin treatment significantly reduced MYC protein half-life, by 1.8-fold, from 41 minutes to 23 minutes with cycloheximide-chase assay (Figure 1 f-g). These results demonstrate that proscillaridin preferentially targets leukemic cells expressing high levels of MYC causing its rapid degradation.

### Proscillaridin Efficiently Targets MYC-Driven Leukemic Stem Cell Populations

We sought to determine whether proscillaridin could target leukemic stem cells since MYC is driving their proliferation (LSCs) (37-39). To explore this possibility, we used two LSC models, a mouse model of T-ALL and a LSC model of human acute myeloid leukemia (AML) (37,40,41). First, pre-LSCs T-ALL cells were isolated from a transgenic mouse model that closely reproduces human T-ALL (42). We previously showed that these pre-LSCs are driven by the *SCL/TAL1* and *LMO1* oncogenes, which depend on *NOTCH1-MYC* pathways, and are resistant to chemotherapeutic drugs used against leukemia (doxorubicin, camptothecin and dexamethasone) (5,37). Proscillaridin (3-10 nM) significantly decreased pre-LSC T-ALL viability by 70% after 4 days of treatment (Figure 2a). Despite being resistant to chemotherapeutic drugs, these pre-LSCs (T-ALL) were sensitive to proscillaridin at clinically relevant doses (37).

**Fig. 2.**
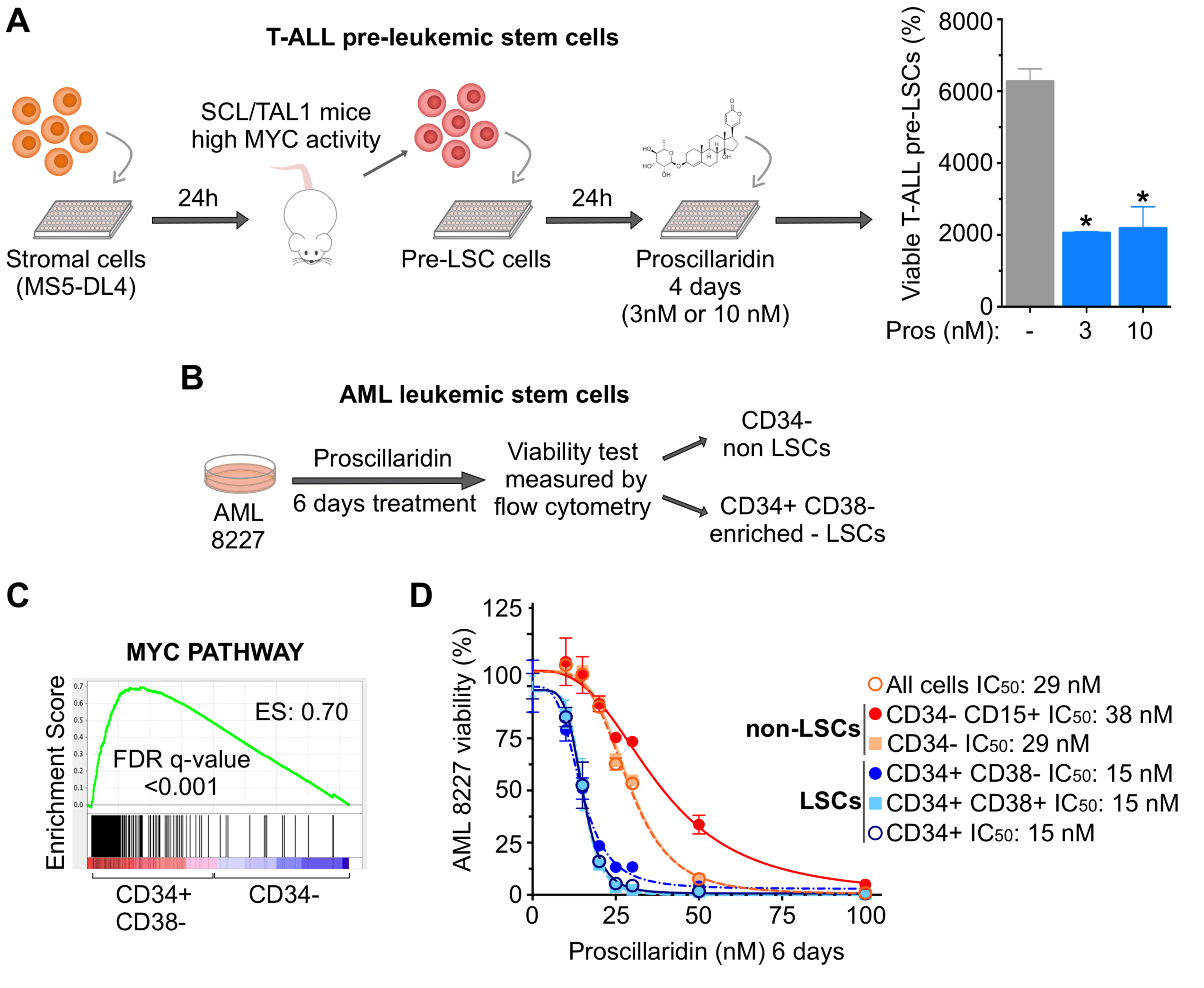
Proscillaridin Targets Leukemic Stem Cells (LSCs). **a** Cell viability assay of T-ALL pre-LSC co-cultured with MS5-DL4 cells. Proscillaridin (3 nM or 10 nM) was added 24h after co-culture, and cells were sorted for pre-LSC viability 4-days post treatment (*indicates P<0.05; ANOVA; n≤3). **b** Cell viability assay of AML 8227 population composed of LSCs (CD34^+^) and non-LSCs (CD34^-^/CD15^+/-^). AML 8227 were treated with proscillaridin (10 nM to 100 nM) for 6 days and cell viability was measured for each cell subgroup by flow cytometry. **c** Gene set enrichment analysis of MYC pathway between two AML 8227 subgroups: LSC-enriched population CD34^+^/CD38^-^ compared to non-LSC population CD34^-^. Enrichment score (ES) and false discovery rate (FDR) rates are shown on the graph. **d** Dose response curves and IC_50_ values after a 6-day proscillaridin treatment (ranging from 10 nM to 100 nM) in each AML 8227 subgroup (n=3).

Then, we used primary human AML 8227 cells, which contain functional LSCs within the CD34^+^ sub-population, and non-LSC cells characterized by CD34^-^ with or without CD15^+^ expression (Figure 2b) (40,41). Gene set enrichment analysis from transcriptomic data published by Lechman et al. revealed that the LSCs-enriched fraction (CD34^+^/CD38^-^) in AML 8227 are enriched for MYC target genes expression as compared to non-LSCs (CD34^-^) (Figure 2c) (40). After 6 days of proscillaridin treatment, bulk AML 8227 cells had an IC_50_ of 29 nM (Figure 2d). Likewise, CD34^-^ with or without CD15^+^ non-LSC cells had IC_50_ of 38 nM and 29 nM, respectively. In contrast, all LSC-enriched AML cells (CD34^+^, CD34^+^/CD38^+^ and CD34^+^/CD38^-^) were more sensitive to proscillaridin with IC_50_ values of 15 nM than non-LSC populations. Altogether, proscillaridin efficiently targets LSC-enriched populations, in both T-ALL and AML models marked by high MYC expression, further supporting its repurposing against MYC-dependent leukemia.

### Proscillaridin Downregulates Cell Proliferation Programs and Induces T-Cell Differentiation

To gain insight into proscillaridin effects against MYC-driven leukemic cells, we investigated drug-induced gene expression changes in T-ALL cells (MOLT-4). By quantitative RT-PCR (qPCR), we found that proscillaridin significantly downregulated MYC mRNA after 16h treatment and up to 90% after 48h (Figure 3a). Then, we used RNA-sequencing to explore transcriptomic effects of proscillaridin (5 nM; 48h) in MOLT-4 cells. After drug treatment, transcriptome analysis showed a downregulation of 2,759 genes (log_2_FC < 0.5; P-value adjusted < 0.05) and concomitant upregulation of 3,271 genes (log_2_FC > 1; P-value adjusted < 0.05; Supplementary Figures 2a and b). Using Metascape, gene ontology analysis revealed that downregulated genes were involved in DNA replication, biosynthesis and metabolic processes (Figure 3b; Supplementary Figure 2c). Consistent with qPCR results, *MYC* transcript was significantly downregulated in our RNA-sequencing data set (Figure 3c). Gene Set Enrichment Analysis showed that MYC PATHWAY (which includes 30 MYC target genes) was significantly downregulated (Figure 3c). Notably, these transcriptomic effects correlated with a 25% decrease of S-phase cells as measured by BrdU staining (Figure 3d; Supplemental Figure 2d). Proscillaridin also significantly downregulated 11 T-cell leukemia master transcription factors (Figure 3e) (43-45). These data support that proscillaridin efficiently inhibits proliferation programs in MYC-driven leukemia.

**Fig. 3.**
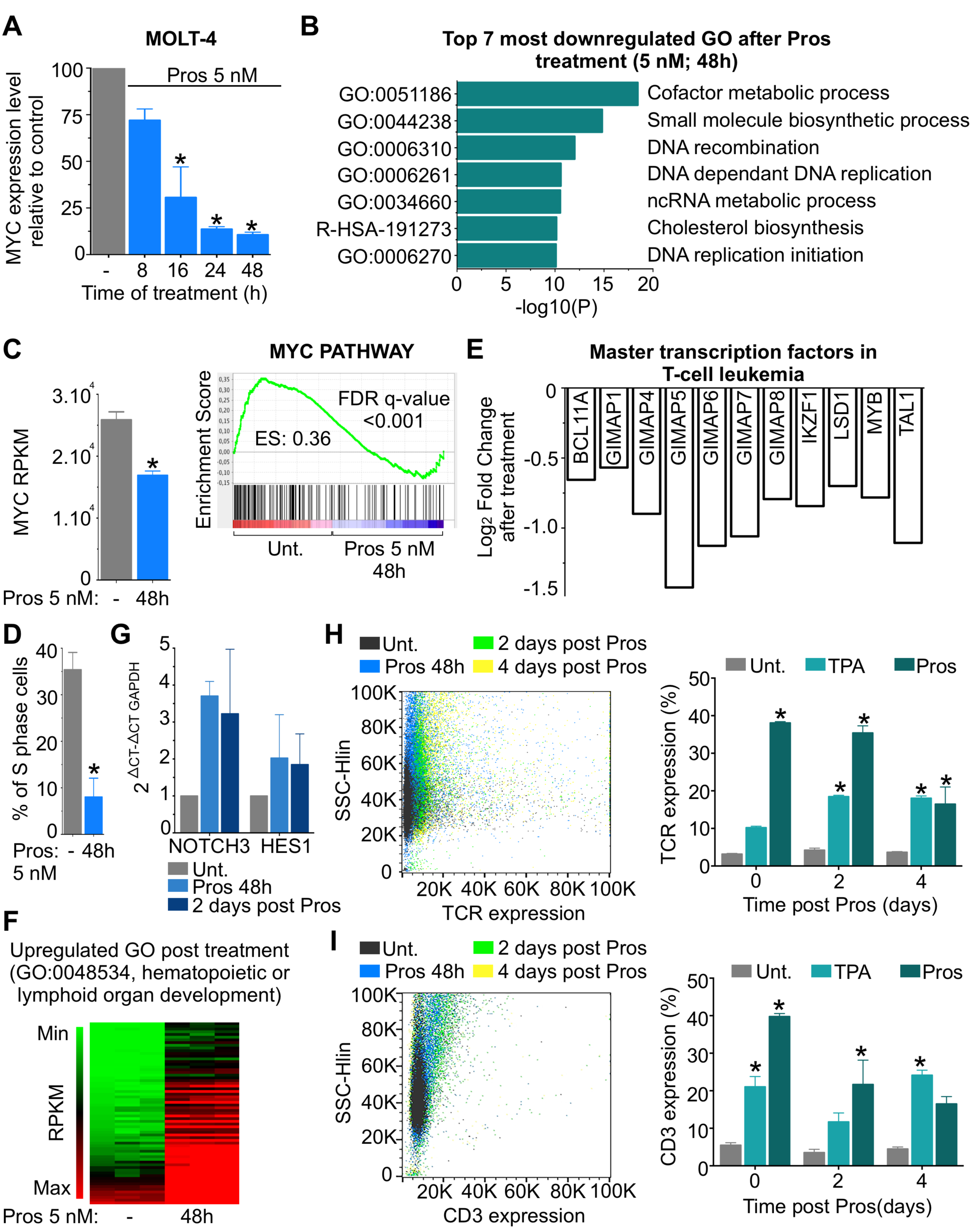
Transcriptomic Profiles from Replicative To Differentiated Phenotype After Low Dose Proscillaridin Treatment in High MYC Expressing Leukemic Cells. **a** Quantitative PCR (qPCR) analysis of *MYC* mRNA expression after proscillaridin treatment (5 nM; 8h to 48h) in MOLT-4 cells, relative to untreated cells and normalized to β-2-microglobulin (t-test: P<0.05; n=3). **b** Transcriptomic analysis by RNA-Sequencing of untreated and proscillaridin-treated (5 nM; 48h) MOLT-4 cells (n = 3). Genes downregulated by proscillaridin treatment (Log_2_ FC<-0.5) were analyzed by Metascape and the top 7 Gene Ontology (GO) pathways are displayed. **c** Left panel, MYC transcript (RPKM) expression after proscillaridin treatment (5 nM; 48h) in RNA-sequencing data set (*indicates P<0.001, t-test, n=3). Right panel, gene set enrichment analysis of MYC pathway before and after proscillaridin treatment (5 nM; 48h) in MOLT-4 cells. Enrichment score (ES) and false discovery rate (FDR) rates are shown on the graph. **d** Effect of proscillaridin treatment (5 nM; 48h) on the percentage of S phase cell population on MOLT-4 cells (* indicates P<0.017, t-test, n = 3). **e** Gene expression values (Log_2_ fold change) obtained from RNA-sequencing in 11 genes downregulated after proscillaridin treatment (5 nM; 48h) in MOLT-4 cells. These genes were selected due to their role as master transcription factors associated in T-cell leukemia. **f** Heat map of gene expression levels (RPKM) involved in differentiation pathways (MOLT-4 cells) before and after proscillaridin treatment (5nM; 48h). **g** Quantitative PCR (qPCR) analysis of *NOTCH3* and *HES1* mRNA expression measured after proscillaridin treatment (5 nM; 48h) and 2 days post treatment, relative to untreated cells and normalized to GAPDH in MOLT-4 cells (n=2). **h** and **i** Left panel, T-cell differentiation markers TCR and CD3 are measured by flow cytometry in MOLT-4 cells after proscillaridin treatment (5 nM; 48h), as well as 2 and 4 days post treatment (n=3). Right panel, percentage of TCR and CD3 expression in MOLT-4 cells. TPA treatment (10 nM; 48h, followed by a 24h resting period) was used as positive control of T-cell differentiation (*indicates P<0.05; ANOVA; n=3).

Gene ontology analysis also revealed that upregulated genes were enriched for hematopoietic or lymphoid organ development, suggesting the onset of leukemia differentiation (Figure 3f and Supplemental Figure 2e). To probe the functional significance of this change, we measured T-cell differentiation markers in MOLT-4 cells before and after treatment. By qPCR, mRNA levels of T-cell differentiation markers *NOTCH3* and its target *HES1* were upregulated after 48h treatment and remained expressed for 2 days after drug removal (Figure 3g) (46,47). By flow cytometry, we measured a significant increase in TCR and CD3 expression, which lasted up to 4 days after drug removal, suggesting the onset of normal T-cell activation. Upregulation of these differentiation markers were in the same range than the levels measured after TPA treatment, a well-known inducer of leukemia differentiation (Figures 3h and i) (48,49). Altogether, proscillaridin treatment produced a transcriptomic shift from a proliferative program to the induction of T-cell leukemia differentiation.

### Proscillaridin Induces Global Loss of Histone H3 Acetylation

Since proscillaridin induced gene expression and phenotypic changes, we hypothesized that it triggers epigenetic effects in high MYC-driven leukemia. We analyzed histone H3 and H4 post-translational modifications by western blotting after 16h to 96h of proscillaridin treatment (5 nM). We found that proscillaridin produced a significant time-dependent reduction (by 75%) of lysine acetylation at H3K9, H3K14, H3K18, H3K27 residues and global loss of H3 acetylation in MOLT-4 cells (Figure 4a; Supplemental Figure 3a). The dramatic reduction in H3K27ac level was confirmed by chromatin immunoprecipitation where H3K27ac antibody pulled-down similar levels of DNA than IgG after treatment (Supplemental Figure 3b). Similar results were obtained in NALM-6 cells after proscillaridin treatment (Supplemental Figure 3c). No change was detected on H4 acetylation or H3 methylation marks (Supplemental Figures 4a and b; and data not shown). Interestingly, loss of H3 acetylation induced global chromatin reorganization in MOLT-4 cells after treatment, as shown by DAPI staining (Supplemental Figure 4c).

**Fig. 4.**
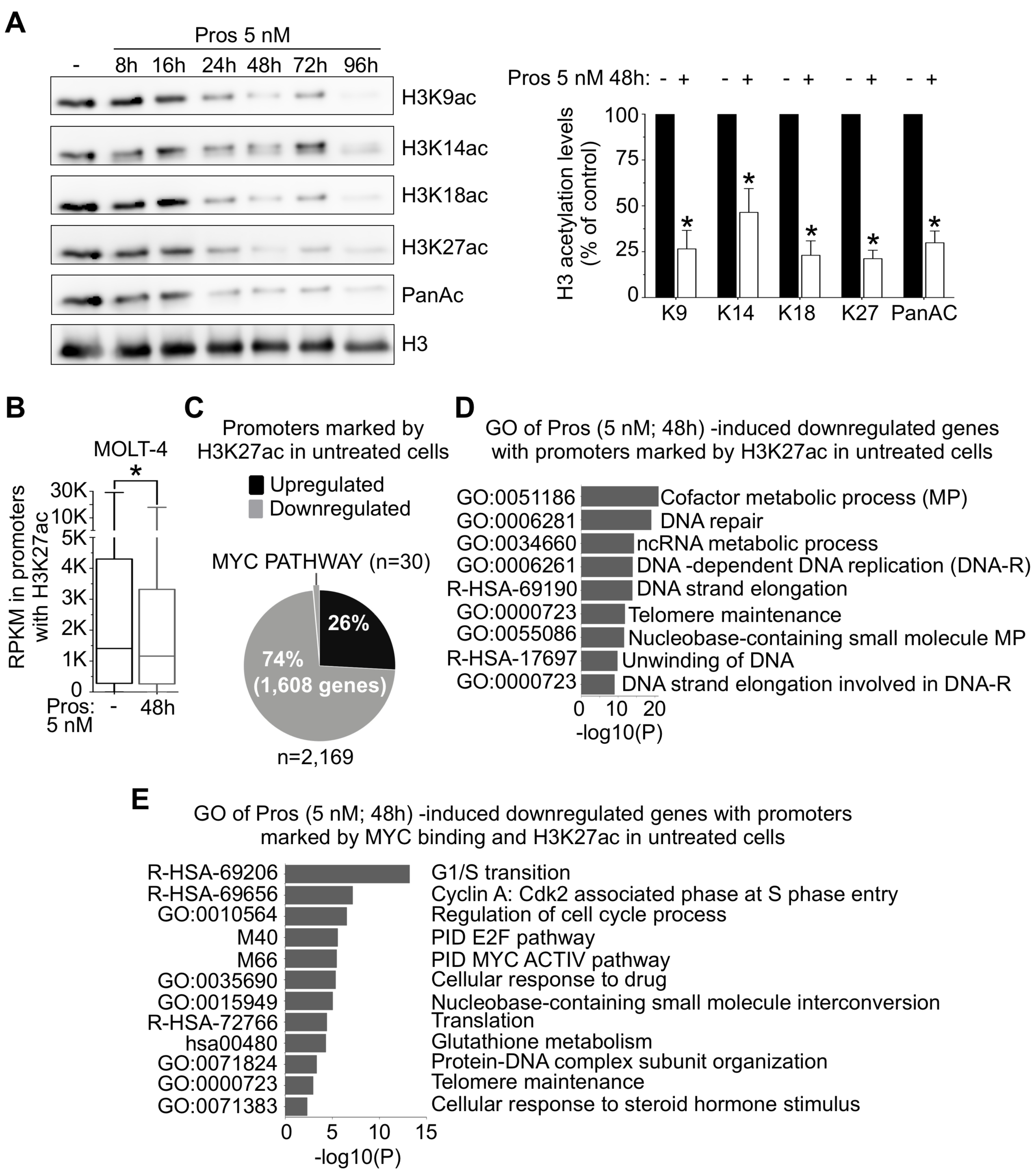
Gene Reprogramming Induced by Proscillaridin Is Associated with Global Acetylation Loss in Histone H3. **a** Right panel, MOLT-4 cells were treated with proscillaridin (5 nM) and histones were acid-extracted after 8, 16, 24, 48, 72 and 96 hours. Histone 3 acetylation levels were assessed using antibodies against K9ac, K14ac, K18ac, K27ac, and total histone 3 acetylation. H3 was used as loading control. Left panel, H3 acetylation levels at 48h treatment were quantified and expressed as a percentage of untreated cells (* indicates P<0.05; ANOVA; n = 3). **b** 2,169 genes are marked by H3K27ac in promoter regions (−500/+500 bp) in untreated MOLT-4 cells. RPKM values of differentially expressed genes (FC > 1; FC<-0.5) from RNA-sequencing data before and after proscillaridin treatment are displayed (* indicates P<0.05; t-test; n = 3). **c** Pie chart shows percentage of upregulated (black) and downregulated (grey, including 30 MYC target) genes marked by H3K27ac in promoters of untreated MOLT-4 cells (5 nM, 48h). **d** Metascape analysis of genes marked by H3K27ac in promoters in untreated MOLT-4 cells and downregulated after proscillaridin treatment (5 nM; 48h). Top 9 GO pathways are displayed. **e** Metascape analysis of the 30 MYC target genes.

We next asked if there was a correlation between loss of H3 acetylation and gene expression changes after proscillaridin treatment. To address this question, we combined our RNA-Seq data pre- and post-treatment with publicly available H3K27ac ChIP-seq data of untreated MOLT-4 (50) since this mark is associated with transcribed regions and is globally lost after treatment (Figure 4a) (23). Among 7,097 genes marked at their promoters with H3K27ac (−500 to +500 bp) in untreated MOLT-4 cells, 2,169 genes were differentially expressed after proscillaridin treatment. Seventy-four percent of those (1,608 genes), marked by H3K27ac in untreated cells, were significantly downregulated after treatment (Figure 4b), which is consistent with the loss of this active epigenetic mark (Figure 4c). Gene ontology analysis of these 1,608 downregulated genes showed a significant relationship with metabolism and proliferation processes (Figure 4d; Supplemental Figure 5a). Among these genes, all MYC pathway genes (n=30) previously described (Figure 3c) were marked by H3K27ac in untreated MOLT-4 cells and were all downregulated by treatment (Figures 4 c and e). Network analysis showed that these MYC target genes are co-expressed simultaneously, and are known to exhibit protein-protein interactions with MYC, confirming the global effect of proscillaridin on MYC pathway (Supplemental Figures 6a and b). By contrast, upregulated genes marked by H3K27ac in untreated cells were associated with apoptosis, negative regulation of proliferation and cell differentiation (Supplemental Figure 5b), corroborating our transcriptomic and functional analyses. Collectively, these results demonstrate that proscillaridin produces global loss of H3 acetylation, which was associated with silencing of genes involved in proliferation and MYC pathway.

**Fig. 5.**
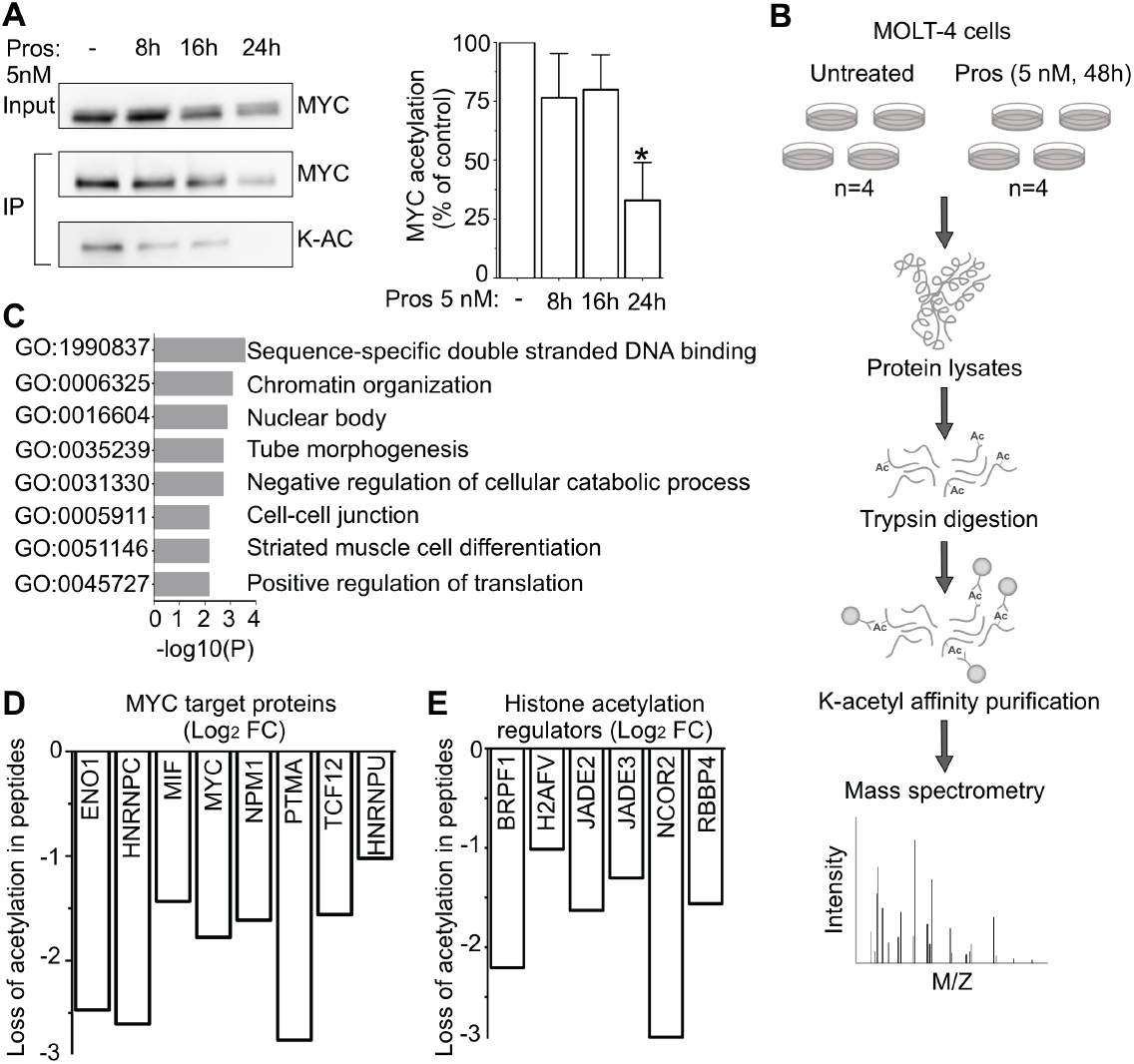
Acetylation Decrease In MYC Targets And Chromatin Regulators Induced by Proscillaridin In High MYC Expressing Cells. **a** Right panel, immunoprecipitation (IP) of MYC total lysine acetylation (K-AC) after proscillaridin treatment (5 nM; 8h-16h-24h) in MOLT-4 cells. Left panel, total lysine acetylation level on MYC was quantified and expressed as a percentage of untreated cells (* indicates P<0.05; ANOVA; n = 3). **b** Lysine acetylome profiling of MOLT-4 cells before and after proscillaridin treatment (5 nM; 48h). **c** Lysine acetylome metascape analysis in 28 peptides with significant loss of acetylation (Log_2_FC<-1) after proscillaridin treatment (5 nM; 48h) in MOLT-4 cells. **d** Log_2_FC of acetylation levels in MYC target proteins (untreated VS treated). **e** Log_2_FC of acetylation levels of histone regulators (untreated VS treated).

**Fig. 6.**
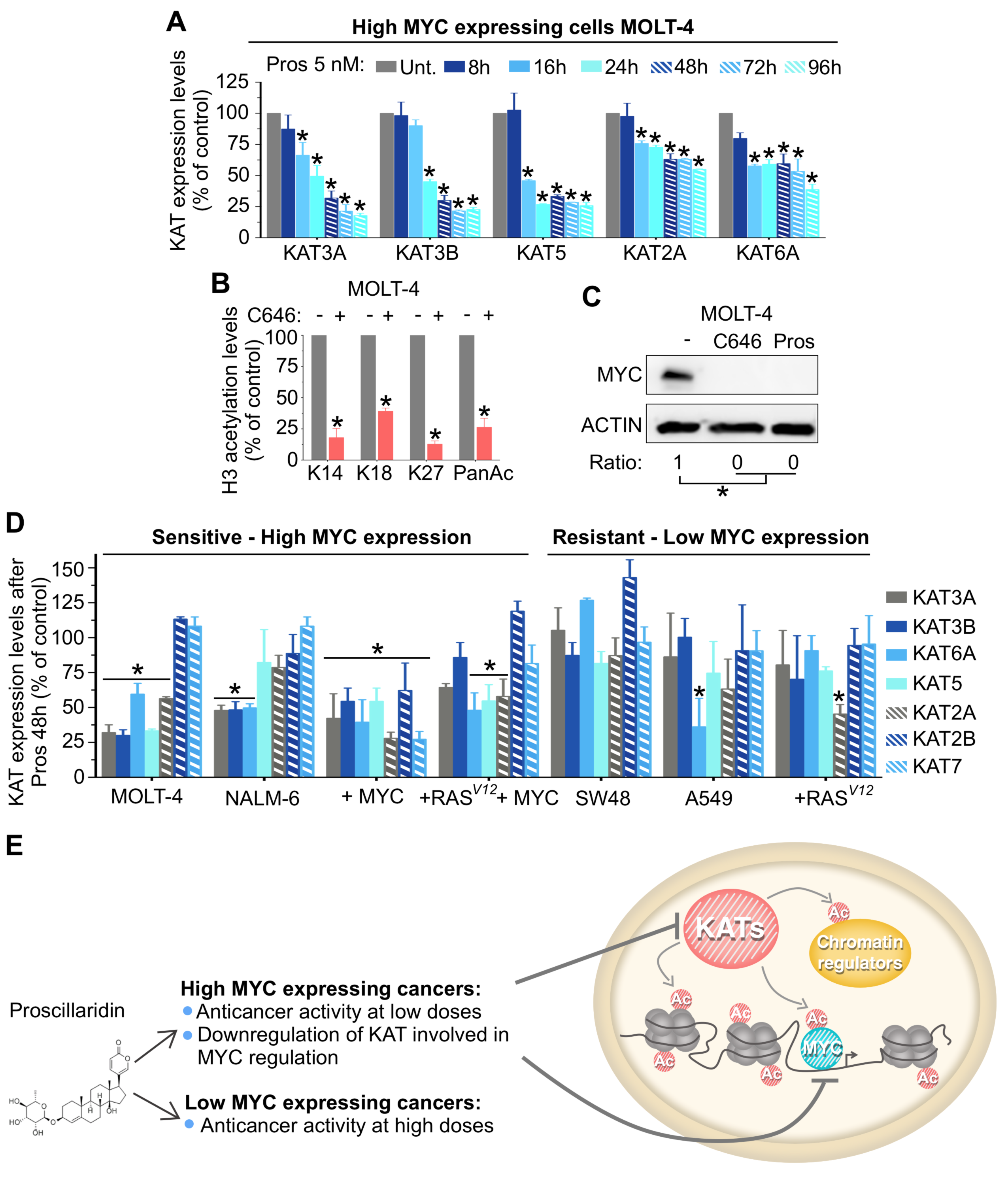
Proscillaridin Treatment Induces Downregulation of KATs Involved In MYC Acetylation. **a** KAT2A (GCN5), KAT3A (CBP), KAT3B (P300), KAT5 (TIP60), and KAT6A (MOZ) expression levels after proscillaridin treatment (5 nM, 48h) were assessed by western blotting in MOLT-4 cells (ACTIN was used as loading control). **b** Quantification of H3 acetylation levels after treatment with KAT3B/A inhibitor C646 (10 µM, 48h) (* indicates P<0.05; t-test; n=3). **c** MYC protein expression after C646 treatment and proscillaridin treatment (5 nM; 48h). ACTIN was used as loading control. **d** MOLT-4, NALM-6, SW48 and A549 cell lines were treated with proscillaridin (5 nM, 48h) and fibroblasts transfected with *RAS*^*V12*^, *MYC* and *RAS*^*V12*^/*MYC* were treated with proscillaridin (70 nM, 48h). KAT3A (CBP), KAT3B (P300), KAT5 (TIP60), KAT2A (GCN5), KAT2B (PCAF), KAT6A (MOZ) and KAT7 (HBO1) expression levels were assessed by western blotting, quantified and expressed as percentage of untreated cells (* indicates P<0.05; t-test; n=3). **e** Scheme showing that proscillaridin targets high MYC expressing leukemic cells by inhibiting the histone acetyltransferases involved in MYC acetylation and stability.

### Proscillaridin Induces Loss of Lysine Acetylation in MYC Target Genes and Chromatin Regulators

We then asked whether depletion of lysine acetylation was extended to non-histone proteins after treatment. First, we measured MYC acetylation levels after 8, 16 and 24h of proscillaridin treatment (5 nM) in MOLT-4 cells, since this posttranslational modification plays a role in its stability (17,51). After MYC immunoprecipitation and probing with a pan-acetyl antibody, we measured a significant time dependent decrease (up to 75%) of MYC total acetylation (Figure 5a).

To further characterize the extent of acetylation loss, we conducted an acetylome study by mass spectrometry on untreated and proscillaridin-treated (5 nM; 48h) MOLT-4 cells (Figure 5b). Two distinct MYC peptides showed a significant reduction in lysine acetylation after treatment, which validated our immunoprecipitation results (Supplemental Figure 7a). Mass spectrometry analysis showed that 28 peptides (including MYC) had a significant loss of lysine acetylation after treatment. Gene ontology of these peptides showed an association with DNA binding and chromatin organization (Figure 5c). Networks analysis showed that these 28 proteins are generally co-expressed, suggesting a connection between their acetylation and expression levels (Supplemental Figure 7b). Interestingly, 8 out of 28 peptides were MYC target proteins, including MYC itself and 6 out of 28 are involved in histone acetylation regulation (Figures 5d and e) (52). Altogether, proscillaridin reduces lysine acetylation of MYC, proteins involved in MYC pathway and several histone acetylation regulators.

### Proscillaridin Downregulates Histone Acetyltransferases Involved in MYC Acetylation more efficiently in high MYC expressing leukemic cells

We next investigated whether acetylation loss was due to a deregulation of histone acetyltransferases (KATs). We measured, by western blotting, KAT levels before and after proscillaridin treatment (5 nM; 8h-96h) in MOLT-4 cells. Proscillaridin produced a time-dependent reduction (up to 80%) of several KATs including KAT3A (CBP), KAT3B (P300), KAT5 (TIP60), KAT2A (GCN5) and KAT6A (MOZ) (Figure 6a, Supplemental Figure 8a). Protein expression levels of KAT2B (PCAF) and KAT7 (HBO1) were not altered by the treatment (Supplemental Figures 8a and b). No significant changes were observed in class I HDACs expression, suggesting that acetylation loss mainly involved KATs downregulation (data not shown). Interestingly, KAT downregulation was observed only at the protein level, since their mRNA levels were not altered after treatment (data not shown). Significant reduction in KAT protein expression including KAT2A, KAT3A, KAT3B and KAT6A, which target histone H3, occurred 8h before significant H3 acetylation loss described previously (Figure 4a). Despite KAT5 decrease, a KAT known to acetylate both H3 and H4, no changes in H4 acetylation (total or on specific lysines) were measured after treatment (Supplemental Figure 4a) (53-57). These results can be explained by the fact that KAT7 (HBO1) expression, which is also involved in H4 acetylation, was not affected by the treatment (58,59). To confirm the effects of KAT downregulation in MOLT-4 cells, we used KAT3A/B pharmacological inhibitor C646. Similar to proscillaridin treatment, C646 (10 µM; 48h) significantly reduced lysine acetylation (H3K14, H3K18, H3K27, and total H3-acetylation), depleted KATs (KAT3A, and KAT3B) and MYC protein levels (Figures 6b and c; Supplemental Figures 8c and d).

Since KATs have overlapping enzymatic activities, we asked if the extent KAT protein downregulation was associated with proscillaridin sensitivity. We compared KAT protein levels before and after proscillaridin treatment in *MYC* overexpressing cancer cells (MOLT-4, NALM-6, *MYC* and *RAS*^*V12*^*+MYC* transformed fibroblasts) versus low *MYC* expressing cancer cells (SW48, A549, and *RAS*^*V12*^ transformed fibroblasts; Figure 6d; Supplemental Figure 9a). Cancer cell lines were treated at 5 nM for 48h, which was clinically relevant and close the IC_50_ values of leukemic cells. Transformed fibroblasts were treated at 70 nM for 48h, which was the IC_50_ value of *MYC* transfected fibroblasts as described in Figure 1b. After treatment, we observed that KAT protein downregulation was more pronounced in drug-sensitive cells with high *MYC* expression as compared to drug-resistant cells with low *MYC* expression. Indeed, proscillaridin induced a significant downregulation of 4/7 KATs in MOLT-4 cells, 3/7 KATs in NALM-6 cells, 3/7 KATs in *RAS*^*V12*^*+MYC* transformed fibroblasts, and 7/7 KATs in *MYC* transformed fibroblasts. Interestingly, these KATs (KAT2A/GCN5, KAT3A/CBP, KAT3B/P300, KAT5/TIP60 and KAT6A/MOZ) are known to acetylate MYC and increase its stability (17,51,60-64). Loss of this specific subset of KATs correlates with loss of MYC acetylation, reduction of MYC half-life and expression, as described previously (Figure 1 and Figure 5). In stark contrast, proscillaridin failed to downregulate more than one KATs in low *MYC* expressing cancer cells (SW48, A549 and *RAS*^*V12*^ transformed fibroblasts). Thus, proscillaridin-induced KAT proteins downregulation occurred more prominently in high *MYC* expressing cells and was associated with clinically relevant IC_50_ values.

Similar analysis was performed on histone H3 acetylation between high *MYC* expressing cancer cells versus low *MYC* expressing cancer cells (Supplemental Figures 9b and c). Proscillaridin induced a significant loss of H3 acetylation in drug-sensitive and *MYC* overexpressing cells (MOLT-4, NALM-6, *MYC* and *RAS*^*V12*^*+MYC* transformed fibroblasts). In proscillaridin-resistant and low expressing *MYC* cancer cells, histone acetylation levels were unchanged in SW48 cells after treatment, which correlated with our previous report (Supplemental Figures 9b and c) (22). By contrast, A549 cells lost significantly H3 acetylation after treatment while *RAS*^*V12*^ transformed fibroblasts lost acetylation fewer lysine residues (K9, K27 and pan-acetyl) (Supplemental Figures 9b-c). These data suggest that proscillaridin sensitivity depends on simultaneous loss of KATs expression involved in the regulation of MYC oncogenic signaling.

### Discussion

The repurposing potential of cardiac glycosides in oncology has been suggested several decades ago and is currently under intense clinical investigation either alone (in prostate cancers, NCT01162135; breast cancer, NCT01763931; and sarcoma, NCT00017446) or in combination with chemotherapy (digoxin with cisplatin in head and neck cancers, NCT02906800; or with epigenetic drug decitabine, NCT03113071) (65,66). Here, our data provide a strong rationale to repurpose cardiac glycoside proscillaridin as a precision medicine in overexpressing MYC leukemia,.

Proscillaridin induced a rapid loss of MYC protein expression in *MYC*-driven leukemia cells and in *MYC*-dependent leukemic stem cells, indicating the potential of controlling leukemia self-renewal capacity. We demonstrated that proscillaridin targeted MYC overexpressing leukemic cells by downregulating KATs involved specifically in MYC acetylation, protein stability and transcriptional program (KAT2A, KAT3A, KAT3B, KAT5 and KAT6A) (17,51,60-64). Importantly, KATs have overlapping activity and targets, suggesting the relevance of targeting these enzymes simultaneously to efficiently reduce acetylation of MYC and its partners (17,61,62,64). Indeed, proscillaridin-induced downregulation of several KATs was observed in proscillaridin-sensitive and MYC overexpressing leukemic cells whereas this effect was sporadic in resistant and low MYC expressing cells. Proscillaridin treatment induced a significant loss of H3 acetylation levels in MYC overexpressing cells whereas loss of histone acetylation was not observed in SW48 cells but was significantly reduced in A549 and in two lysine residues in *RAS*^*v*12^-transfected fibroblasts, suggesting that loss of H3 acetylation is not sufficient to alter cell viability in low MYC expressing cells. These data highlight the importance of acetylation levels in non-histone proteins as a potential therapeutic target in MYC overexpressing leukemia. Experiments are ongoing to address this specific question.

Lysine acetylation is a dynamic process that can be modulated within minutes and it is regulated in histone and non-histone proteins by the redundant activity of KATs. Therefore, cancer cells may rapidly recover from incomplete pharmacological inhibition or from the specific inhibition of a particular KAT (53). Here, we showed that proscillaridin treatment in MYC overexpressing leukemia cells, led to the downregulation of several KATs, produced MYC inhibition, and induced persistent leukemia cell differentiation, which was maintained for several days after drug removal. Therefore, this study supports a strategy of simultaneously targeting several KATs to reduce efficiently acetylation in histone and in non-histone proteins, which overcome the redundant activity of KATs. The mechanism implicated in proscillaridin-induced KATs downregulation is under investigation. Overall, we conclude that cardiac glycoside proscillaridin efficiently targets MYC overexpressing leukemia by reducing global lysine acetylation leading to MYC pathway inhibition and the induction of leukemia differentiation (Figure 6e).

## Supporting information

## Acknowledgements

We thank Catherine Legros for the graphical help, and Dr André Tremblay, Dr Audrey Claing, Dr Elie Haddad and Dr Carolina Alfieri for providing cell lines and materials.

## Author contributions

E.M.D.C. and N.J.M.R. designed the study and wrote the manuscript. E.M.D.C., G.A., G. McI., A.B., V. B-L., C.R., M. R., M. B., K. E., EE. F-D., A. H., T. H., C. B., and S. McG. performed the experiments. C.R., M.C., P.S.O and D.S performed RNA sequencing experiments and bioinformatics analyses. J.R. J., N. K., Y. S., M. D. performed mass spectrometry experiments and acetylome studies. The authors declare no conflict of interest.

## References

1. Bradner JE, Hnisz D, Young RA. Transcriptional Addiction in Cancer. Cell 2017;168(4):629–43 doi 10.1016/j.cell.2016.12.013.

2. Fernandez PC, Frank SR, Wang L, Schroeder M, Liu S, Greene J, et al. Genomic targets of the human c-Myc protein. Genes Dev 2003;17(9):1115–29 doi 10.1101/gad.1067003.

3. Dang CV. MYC on the path to cancer. Cell 2012;149(1):22–35 doi 10.1016/j.cell.2012.03.003.

4. Delgado MD, Leon J. Myc roles in hematopoiesis and leukemia. Genes Cancer 2010;1(6):605–16 doi 10.1177/1947601910377495.

5. Gerby B, Tremblay CS, Tremblay M, Rojas-Sutterlin S, Herblot S, Hebert J, et al. SCL, LMO1 and Notch1 reprogram thymocytes into self-renewing cells. PLoS Genet 2014;10(12):e1004768 doi 10.1371/journal.pgen.1004768.

6. Felsher DW, Bishop JM. Reversible tumorigenesis by MYC in hematopoietic lineages. Molecular cell 1999;4(2):199–207.

7. Marinkovic D, Marinkovic T, Mahr B, Hess J, Wirth T. Reversible lymphomagenesis in conditionally c-MYC expressing mice. International journal of cancer Journal international du cancer 2004;110(3):336–42 doi 10.1002/ijc.20099.

8. Koh CM, Sabo A, Guccione E. Targeting MYC in cancer therapy: RNA processing offers new opportunities. Bioessays 2016;38(3):266–75 doi 10.1002/bies.201500134.

9. Posternak V, Cole MD. Strategically targeting MYC in cancer. F1000Res 2016;5 doi 10.12688/f1000research.7879.1.

10. Filippakopoulos P, Qi J, Picaud S, Shen Y, Smith WB, Fedorov O, et al. Selective inhibition of BET bromodomains. Nature 2010;468(7327):1067–73 doi 10.1038/nature09504.

11. Delmore JE, Issa GC, Lemieux ME, Rahl PB, Shi J, Jacobs HM, et al. BET bromodomain inhibition as a therapeutic strategy to target c-Myc. Cell 2011;146(6):904–17 doi 10.1016/j.cell.2011.08.017.

12. Roe JS, Mercan F, Rivera K, Pappin DJ, Vakoc CR. BET Bromodomain Inhibition Suppresses the Function of Hematopoietic Transcription Factors in Acute Myeloid Leukemia. Molecular cell 2015;58(6):1028–39 doi 10.1016/j.molcel.2015.04.011.

13. Perez-Salvia M, Esteller M. Bromodomain inhibitors and cancer therapy: From structures to applications. Epigenetics: official journal of the DNA Methylation Society 2017;12(5):323–39 doi 10.1080/15592294.2016.1265710.

14. Shu S, Lin CY, He HH, Witwicki RM, Tabassum DP, Roberts JM, et al. Response and resistance to BET bromodomain inhibitors in triple-negative breast cancer. Nature 2016;529(7586):413–7 doi 10.1038/nature16508.

15. Fong CY, Gilan O, Lam EY, Rubin AF, Ftouni S, Tyler D, et al. BET inhibitor resistance emerges from leukaemia stem cells. Nature 2015;525(7570):538–42 doi 10.1038/nature14888.

16. Kurimchak AM, Shelton C, Duncan KE, Johnson KJ, Brown J, O’Brien S, et al. Resistance to BET Bromodomain Inhibitors Is Mediated by Kinome Reprogramming in Ovarian Cancer. Cell Rep 2016;16(5):1273–86 doi 10.1016/j.celrep.2016.06.091.

17. Faiola F, Liu X, Lo S, Pan S, Zhang K, Lymar E, et al. Dual regulation of c-Myc by p300 via acetylation-dependent control of Myc protein turnover and coactivation of Myc-induced transcription. Mol Cell Biol 2005;25(23):10220–34 doi 10.1128/MCB.25.23.10220-10234.2005.

18. Glozak MA, Sengupta N, Zhang X, Seto E. Acetylation and deacetylation of non-histone proteins. Gene 2005;363:15–23 doi 10.1016/j.gene.2005.09.010.

19. Luebben WR, Sharma N, Nyborg JK. Nucleosome eviction and activated transcription require p300 acetylation of histone H3 lysine 14. Proceedings of the National Academy of Sciences of the United States of America 2010;107(45):19254–9 doi 10.1073/pnas.1009650107.

20. Dahlin JL, Nelson KM, Strasser JM, Barsyte-Lovejoy D, Szewczyk MM, Organ S, et al. Assay interference and off-target liabilities of reported histone acetyltransferase inhibitors. Nature communications 2017;8(1):1527 doi 10.1038/s41467-017-01657-3.

21. Raynal NJ, Da Costa EM, Lee JT, Gharibyan V, Ahmed S, Zhang H, et al. Repositioning FDA-Approved Drugs In Combination With Epigenetic Drugs To Reprogram Colon Cancer Epigenome. Molecular Cancer Therapeutics 2017;16(2):397–407 doi 10.1158/1535-7163.

22. Raynal NJ, Lee JT, Wang Y, Beaudry A, Madireddi P, Garriga J, et al. Targeting Calcium Signaling Induces Epigenetic Reactivation of Tumor Suppressor Genes in Cancer. Cancer research 2016;76(6):1494–505 doi 10.1158/0008-5472.CAN-14-2391.

23. Prassas I, Diamandis EP. Novel therapeutic applications of cardiac glycosides. Nature reviews Drug discovery 2008;7(11):926–35 doi 10.1038/nrd2682.

24. Newman RA, Yang P, Pawlus AD, Block KI. Cardiac glycosides as novel cancer therapeutic agents. Mol Interv 2008;8(1):36–49 doi 8/1/36 [pii]10.1124/mi.8.1.8.

25. Jain S, Vaidyanathan B. Digoxin in management of heart failure in children: Should it be continued or relegated to the history books? Ann Pediatr Cardiol 2009;2(2):149–52 doi 10.4103/0974-2069.58317.

26. Tailler M, Senovilla L, Lainey E, Thepot S, Metivier D, Sebert M, et al. Antineoplastic activity of ouabain and pyrithione zinc in acute myeloid leukemia. Oncogene 2012;31(30):3536–46 doi onc2011521 [pii] 10.1038/onc.2011.521.

27. Prassas I, Paliouras M, Datti A, Diamandis EP. High-throughput screening identifies cardiac glycosides as potent inhibitors of human tissue kallikrein expression: implications for cancer therapies. Clin Cancer Res 2008;14(18):5778–84 doi 10.1158/1078-0432.CCR-08-0706.

28. Denicolai E, Baeza-Kallee N, Tchoghandjian A, Carre M, Colin C, Jiglaire CJ, et al. Proscillaridin A is cytotoxic for glioblastoma cell lines and controls tumor xenograft growth in vivo. Oncotarget 2014;5(21):10934–48 doi 10.18632/oncotarget.2541.

29. Kometiani P, Liu L, Askari A. Digitalis-induced signaling by Na+/K+-ATPase in human breast cancer cells. Mol Pharmacol 2005;67(3):929–36 doi 10.1124/mol.104.007302.

30. Dobin A, Davis CA, Schlesinger F, Drenkow J, Zaleski C, Jha S, et al. STAR: ultrafast universal RNA-seq aligner. Bioinformatics 2013;29(1):15–21 doi 10.1093/bioinformatics/bts635.

31. Love MI, Huber W, Anders S. Moderated estimation of fold change and dispersion for RNA-seq data with DESeq2. Genome biology 2014;15(12):550 doi 10.1186/s13059-014-0550-8.

32. Downey M, Johnson JR, Davey NE, Newton BW, Johnson TL, Galaang S, et al. Acetylome profiling reveals overlap in the regulation of diverse processes by sirtuins, gcn5, and esa1. Mol Cell Proteomics 2015;14(1):162–76 doi 10.1074/mcp.M114.043141.

33. Cox J, Mann M. MaxQuant enables high peptide identification rates, individualized p.p.b.-range mass accuracies and proteome-wide protein quantification. Nat Biotechnol 2008;26(12):1367–72 doi 10.1038/nbt.1511.

34. Choi M, Chang CY, Clough T, Broudy D, Killeen T, MacLean B, et al. MSstats: an R package for statistical analysis of quantitative mass spectrometry-based proteomic experiments. Bioinformatics 2014;30(17):2524–6 doi 10.1093/bioinformatics/btu305.

35. Lin CY, Loven J, Rahl PB, Paranal RM, Burge CB, Bradner JE, et al. Transcriptional amplification in tumor cells with elevated c-Myc. Cell 2012;151(1):56–67 doi 10.1016/j.cell.2012.08.026.

36. Gabay M, Li Y, Felsher DW. MYC activation is a hallmark of cancer initiation and maintenance. Cold Spring Harbor perspectives in medicine 2014;4(6) doi 10.1101/cshperspect.a014241.

37. Gerby B, Veiga DFT, Krosl J, Nourreddine S, Ouellette J, Haman A, et al. High-throughput screening in niche-based assay identifies compounds to target preleukemic stem cells. The Journal of Clinical Investigation 2016;126(12):4569–84 doi 10.1172/JCI86489.

38. Shlush LI, Zandi S, Mitchell A, Chen WC, Brandwein JM, Gupta V, et al. Identification of pre-leukaemic haematopoietic stem cells in acute leukaemia. Nature 2014;506(7488):328–33 doi 10.1038/nature13038.

39. Corces-Zimmerman MR, Hong WJ, Weissman IL, Medeiros BC, Majeti R. Preleukemic mutations in human acute myeloid leukemia affect epigenetic regulators and persist in remission. Proc Natl Acad Sci U S A 2014;111(7):2548–53 doi 10.1073/pnas.1324297111.

40. Lechman ER, Gentner B, Ng SW, Schoof EM, van Galen P, Kennedy JA, et al. miR-126 Regulates Distinct Self-Renewal Outcomes in Normal and Malignant Hematopoietic Stem Cells. Cancer Cell 2016;29(2):214–28 doi 10.1016/j.ccell.2015.12.011.

41. Laverdiere I, Boileau M, Neumann AL, Frison H, Mitchell A, Ng SWK, et al. Leukemic stem cell signatures identify novel therapeutics targeting acute myeloid leukemia. Blood Cancer J 2018;8(6):52 doi 10.1038/s41408-018-0087-2.

42. Tremblay M, Tremblay CS, Herblot S, Aplan PD, Hebert J, Perreault C, et al. Modeling T-cell acute lymphoblastic leukemia induced by the SCL and LMO1 oncogenes. Genes & development 2010;24(11):1093–105 doi 10.1101/gad.1897910.

43. Hnisz D, Abraham BJ, Lee TI, Lau A, Saint-Andre V, Sigova AA, et al. Super-enhancers in the control of cell identity and disease. Cell 2013;155(4):934–47 doi 10.1016/j.cell.2013.09.053.

44. Niederriter AR, Varshney A, Parker SC, Martin DM. Super Enhancers in Cancers, Complex Disease, and Developmental Disorders. Genes (Basel) 2015;6(4):1183–200 doi 10.3390/genes6041183.

45. Whyte WA, Orlando DA, Hnisz D, Abraham BJ, Lin CY, Kagey MH, et al. Master transcription factors and mediator establish super-enhancers at key cell identity genes. Cell 2013;153(2):307– 19 doi 10.1016/j.cell.2013.03.035.

46. Bellavia D, Campese AF, Vacca A, Gulino A, Screpanti I. Notch3, another Notch in T cell development. Semin Immunol 2003;15(2):107–12.

47. Tsukumo S, Yasutomo K. Notch governing mature T cell differentiation. J Immunol 2004;173(12):7109–13.

48. Kuhns MS, Davis MM, Garcia KC. Deconstructing the form and function of the TCR/CD3 complex. Immunity 2006;24(2):133–9 doi 10.1016/j.immuni.2006.01.006.

49. Carlsson M, Matsson P, Rosen A, Sundstrom C, Totterman TH, Nilsson K. Phorbol ester and B cell-stimulatory factor synergize to induce B-chronic lymphocytic leukemia cells to simultaneous immunoglobulin secretion and DNA synthesis. Leukemia 1988;2(11):734–44.

50. Abraham BJ, Hnisz D, Weintraub AS, Kwiatkowski N, Li CH, Li Z, et al. Small genomic insertions form enhancers that misregulate oncogenes. Nat Commun 2017;8:14385 doi 10.1038/ncomms14385.

51. Vervoorts J, Luscher-Firzlaff JM, Rottmann S, Lilischkis R, Walsemann G, Dohmann K, et al. Stimulation of c-MYC transcriptional activity and acetylation by recruitment of the cofactor CBP. EMBO Rep 2003;4(5):484–90 doi 10.1038/sj.embor.embor821.

52. Avvakumov N, Lalonde ME, Saksouk N, Paquet E, Glass KC, Landry AJ, et al. Conserved molecular interactions within the HBO1 acetyltransferase complexes regulate cell proliferation. Molecular and cellular biology 2012;32(3):689–703 doi 10.1128/MCB.06455-11.

53. Di Martile M, Del Bufalo D, Trisciuoglio D. The multifaceted role of lysine acetylation in cancer: prognostic biomarker and therapeutic target. Oncotarget 2016;7(34):55789–810 doi 10.18632/oncotarget.10048.

54. Jin Q, Yu LR, Wang L, Zhang Z, Kasper LH, Lee JE, et al. Distinct roles of GCN5/PCAF-mediated H3K9ac and CBP/p300-mediated H3K18/27ac in nuclear receptor transactivation. EMBO J 2011;30(2):249–62 doi 10.1038/emboj.2010.318.

55. Henry RA, Kuo YM, Andrews AJ. Differences in Specificity and Selectivity Between CBP and p300 acetylation of histone H3 and H3/H4. Biochemistry 2013 doi 10.1021/bi400684q.

56. Huang F, Abmayr SM, Workman JL. Regulation of KAT6 Acetyltransferases and Their Roles in Cell Cycle Progression, Stem Cell Maintenance, and Human Disease. Molecular and cellular biology 2016;36(14):1900–7 doi 10.1128/MCB.00055-16.

57. Kimura A, Horikoshi M. Tip60 acetylates six lysines of a specific class in core histones in vitro. Genes Cells 1998;3(12):789–800.

58. Foy RL, Song IY, Chitalia VC, Cohen HT, Saksouk N, Cayrou C, et al. Role of Jade-1 in the histone acetyltransferase (HAT) HBO1 complex. J Biol Chem 2008;283(43):28817–26 doi 10.1074/jbc.M801407200.

59. Avvakumov N, Cote J. The MYST family of histone acetyltransferases and their intimate links to cancer. Oncogene 2007;26(37):5395–407 doi 10.1038/sj.onc.1210608.

60. McMahon SB, Wood MA, Cole MD. The essential cofactor TRRAP recruits the histone acetyltransferase hGCN5 to c-Myc. Molecular and cellular biology 2000;20(2):556–62.

61. Zhang K, Faiola F, Martinez E. Six lysine residues on c-Myc are direct substrates for acetylation by p300. Biochem Biophys Res Commun 2005;336(1):274–80 doi 10.1016/j.bbrc.2005.08.075.

62. Frank SR, Parisi T, Taubert S, Fernandez P, Fuchs M, Chan HM, et al. MYC recruits the TIP60 histone acetyltransferase complex to chromatin. EMBO Rep 2003;4(6):575–80 doi 10.1038/sj.embor.embor861.

63. Sheikh BN, Lee SC, El-Saafin F, Vanyai HK, Hu Y, Pang SH, et al. MOZ regulates B-cell progenitors and, consequently, Moz haploinsufficiency dramatically retards MYC-induced lymphoma development. Blood 2015;125(12):1910–21 doi 10.1182/blood-2014-08-594655.

64. Patel JH, Du Y, Ard PG, Phillips C, Carella B, Chen CJ, et al. The c-MYC oncoprotein is a substrate of the acetyltransferases hGCN5/PCAF and TIP60. Molecular and cellular biology 2004;24(24):10826–34 doi 10.1128/MCB.24.24.10826-10834.2004.

65. Stenkvist B, Bengtsson E, Eriksson O, Holmquist J, Nordin B, Westman-Naeser S. Cardiac glycosides and breast cancer. Lancet 1979;1(8115):563.

66. Stenkvist B, Bengtsson E, Eklund G, Eriksson O, Holmquist J, Nordin B, et al. Evidence of a modifying influence of heart glucosides on the development of breast cancer. Anal Quant Cytol 1980;2(1):49–54.

