## Supplementary material for "Targeting MYC Overexpressing Leukemia with Cardiac Glycoside Proscillaridin Through Downregulation of Histone Acetyltransferases"

**Supplementary Materials and Methods**

Cell Culture and Drug Treatments

MOLT-4 (T-cell acute lymphoblastic leukemia), NALM-6 (pre-B-cell acute lymphoblastic leukemia), REH (pre-B acute lymphoblastic leukemia), KOPN-8 (B-cell precursor acute lymphoblastic leukemia), CCRF-CEM (T-cell acute lymphoblastic leukemia), K562 (chronic myelogenous leukemia) and DLD-1 (colorectal-adenocarcinoma) cells were cultured in RPMI-1640 (Life Technologies, USA) supplemented with 10% fetal bovine serum (FBS, Wisent Inc., Canada). THP-1 (acute monocytic leukemia) cells were cultured with the same media but it was also supplemented with 0.05 mM of 2-mercaptoethanol. CALU-6 (anasplastic carcinoma) cells were cultured in MEM (Life Technologies, USA). RD (embryonal rhabdomyosarcoma) and MCF7 (breast adenocarcinoma) cells were cultured in DMEM (Life Technologies, USA) supplemented with 10% FBS. U2OS (osteosarcoma) cells were cultured in McCoy’s (Life Technologies, United Kingdom) supplemented with 10% FBS. A549 (lung carcinoma) cells were cultured in F12K (GE Healthcare Life Sciences, Canada) supplemented with 10% of FBS. SW48 (colorectal adenocarcinoma) cells were cultured in Leibovitz L-15 supplemented with 10% FBS. Cell lines were purchased from American Type Culture Collection and tested for mycoplasma contamination. All cell lines were culture in a humidified incubator with 5% CO_2_ at 37°C, except SW48 cells that were cultured with 1% CO_2_. K562 cells were provided by Dr Elie Haddad and MCF7 cells were supplied by Dr Audrey Claing. AML 8227 were cultured in StemSpan^TM^ SFEM II (STEMCELL Technologies) supplemented with growth factors (Life Technologies: 10 ng/mL IL3, IL6 and G-CSF, 25 ng/mL TPO, 50 ng/mL SCF and FLT3L, and 100 ng/mL SCF) and penicillin-streptomycin (Life Technologies). Phorbol 12-myristate 13-acetate (TPA) was purchased from Sigma-Aldrich, and C646 (Histone Acetyltransferase KAT3B/KAT3A Inhibitor) was purchased from EMD Millipore.

Growth Inhibition

ORFLO MOXI Mini Automated Cell Counter was used to analyze growth inhibition after 48h treatment. IC_50_ values were calculated with GraphPad Prism software. GUAVA EasyCyte (Millipore Sigma-Aldrich) was used to measure cell viability after 24h treatment. Viacount reagent (Millipore Sigma-Aldrich cat#4000-0040) was added prior to analysis. For AML 8227 cells, proscillaridin was added at specified concentrations and incubated for 6 days. Cells were analyzed by flow cytometry. Phenotype and viability were assessed using CD34-APC (581), CD38-PE (HB-7), CD15-FITC (HI98) and SYTOX Blue (Life Technologies). All antibodies were purchased from BioLegend. Flow cytometry was performed using a LSRFortessa fitted with a high-throughput sampler (BD Biosciences).

Nuclei Imaging

Cell pellets were fixed for 10 minutes at room temperature with 3.7% formaldehyde (Anachemia, cat#41883-360) and then permeabilized for 10 minutes at room temperature with 0.1% triton (PBS). Nuclei were stained and mounted with ProLong™ Diamond Antifade Mountant with DAPI (Invitrogen; cat#36962). Images were acquired using Leica TCS SP8 confocal microscope.

Cell Cycle Analysis

Cells were treated with proscillaridin for 48 hours at 3 different concentrations (2.5, 5, or 7.5 nM). Cells were then harvested, fixed, and stained with bromodeoxyuridine (BrdU) and/or 7-amino-actinomycin (7-AAD) (BD Pharmingen BrdU Flow Kit). Samples were analyzed by Fluorescence-activated Cell Sorting (FACS) on a BD FACS Canto II.

T-cell differentiation assay

GUAVA EasyCyte (Millipore Sigma-Aldrich) was used to measure T-cells differentiation on MOLT-4 cells. TCR expression was measured by staining MOLT-4 cells with APC IGM $\kappa$ Isotype Control (BD pharmigen; cat#550883APC) and TCR alpha beta (BD pharmigen, cat#563826) and IGG. CD3 expression was measured by staining MOLT-4 cells with PE-Cy^tm^ IGG1 $\kappa$ isotype (BD pharmigen, cat#557872) and PE-Cy^tm^ CD3 (BD pharmigen, cat#560910).

Antibodies for western blots

The following primary antibodies were used to detect expression levels of isolated proteins: C-MYC (1:5000 Abcam cat#AB32072), RAS (1:2500 Millipore HA-RAS clone MC57, cat#ng1905692); KAT3A (1:1000 Cell Signaling cat#7389), KAT3B (1:2000 Active Motif cat#61402), KAT2A (1:1000 Santa Cruz cat#sc-365321), KAT2B (1:1000 Cell Signaling cat#3378), KAT5 (1:2500 Abcam cat#ab137518), KAT6A (1:1000 Active Motif cat#39868), KAT7 (1:1000 Bethyl Labs cat#A302-224A-T); and Actin (1:5000 Sigma-Aldrich cat#A2228).

The following primary antibodies were used to detect histone acetylation marks on H3 and H4 subunits: H3K9ac (1:5000 Active Motif cat#39917), H3K14ac (1:5000 Active Motif cat#39698), H3K18ac (1:2500 Active Motif #39588), H3K27ac (1:5000 Active Motif #39134), H3ac (pan-acetyl) (1:5000 Active Motif cat#39140), H3 (1:5000 Active Motif cat#39763), H4K5ac (1:2000 Active Motif cat#39700), H4K8ac (1:5000 Active Motif cat#39172); H4K12ac (1:5000 Active Motif cat#39928), H4K16ac (1:5000 Active Motif cat#39167), H4K20ac (1:2000 Active Motif cat#61531), H4ac (pan-acetyl) (1:5000 Active Motif cat#39244) and H4 (1:5000 Active Motif cat#39270). Densitometric analysis was performed using ImageJ software. ANOVA tests were performed with GraphPad Prism software.

Acetylome analysis by mass spectrometry

Sample preparation: Pellets were resuspended in 1 ml of lysis buffer (8 M Urea, 0.1 M Tris-HCl pH 8.0, 150 mM NaCl, 10 mM sodium butyrate (Sigma 303410), 10 mM nicotinamide (Sigma 3376), with Roche protease inhibitor tablet w/o EDTA and lysed using 5 passes through a 21-gauge needle followed by clarification at 15,000 g for 10 minutes at room temperature. The supernatant was quantitated using a Pierce BCA protein kit. One mg of protein was reduced with TCEP (Sigma C4706–2, 4 mM final concentration), alkylated with iodoacetamide (Sigma L1149–5G, prepared fresh in water at 10 mM), and quenched with 10 mM DTT. Samples were diluted to < 2 M urea with 0.1 M Tris-HCl pH 8.0 prior to overnight digestion with trypsin (Pierce 90058, in 50 mM acetic acid, ~15 µg/sample) at room temperature.

Digested samples were acidified using 10% Trifluoroacetic acid (TFA) to a final pH of < 3. Subsequently, insoluble material was precipitated by centrifugation at 10,000 g at room temperature. Clarified supernatant was loaded onto a Sep Pak tC18 column (Waters) activated with 1 ml 80% ACN/0.1% TFA and equilibrated with 3 × 1 ml 0.1% TFA. The columns were washed with 5 × 1 ml 0.1% TFA and eluted with 1 ml 40% ACN/0.1% TFA. The eluted material was frozen and lyophilized over 48 h to completely remove TFA. Dried peptides were resuspended in 800 µl IP buffer (0.1 M Tris-HCl pH 8, 50 mM NaCl, Roche protease inhibitor tablet w/o EDTA) and clarified by centrifugation for 2 minutes at 10,000 g. For purification of acetylated peptides, supernatant was incubated with 50 µl packed volume of ImmuneChem anti-acetyl-lysine agarose beads (ICP0388). Immunoprecipitation was carried out at 4°C for 2 hours. Beads were washed twice with 1 ml lysis buffer and twice with 1 ml water prior to elution with 0.15% TFA at room temperature. Two elutions of 60 µl (10 minutes each) were pooled for analysis.

Mass spectrometry set up: In preparation for mass spectrometry, samples were loaded onto Pierce C18 spin tips (84850) previously activated with 0.1% TFA/80% acetonitrile (ACN) and washed with 0.1% TFA. Following application of the sample, tips were washed twice with 0.1 % TFA and eluted with 0.1% TFA/40% ACN. The elutions were dried on a speed-vac without heat for 4 hours. Acetyl-lysine-enriched peptides were run in technical duplicate on a Thermo Fisher Orbitrap Fusion mass spectrometry system. This system is equipped with an Easy nLC 1200 ultra-high pressure liquid chromatography system interfaced via a Nanospray Flex nanoelectrospray source. The samples were injected on a C18 reverse phase column (25 cm x 75 um packed with ReprosilPur C18 AQ 1.9 µm particles). An organic gradient from 5% to 30% ACN in 0.1% formic acid over 112 minutes at a flow rate of 300 nl/min was used to separate peptides. Spectra were continuously collected by the MS in a data-dependent manner throughout the gradient, acquiring a full scan in the Orbitrap (at 120,000 resolution with an AGC target of 200,000 and a maximum injection time of 100 ms) followed by as many MS/MS scans as could be acquired on the most abundant ions in 3s in the dual linear ion trap. A rapid scan type with an intensity threshold of 5000, HCD collision energy of 29%, AGC target of 10,000, a maximum injection time of 35 ms, and an isolation width of 1.6 m/z was used. All singly and unassigned charge states were rejected. Dynamic exclusion was enabled with a repeat count of 1. The exclusion duration used was 20 s with an exclusion mass width of +/- 10 ppm.

Modification and determination of changes: Variable modifications were allowed for N-terminal protein acetylation, methionine oxidation, and lysine acetylation. A static modification was indicated for carbamidomethyl cysteine. All other settings were left as default in the MaxQuant program. Decoy hits, contaminants and peptides that did not contain acetyl-lysine residues were removed. All samples were normalized across fractions by median-centering the log2-transformed MS1-intensity distributions. MSstats group comparison function was run with the following options: no interaction terms for missing values, no interference, unequal intensity feature variance, restricted technical and biological scope of replication. Statistically significant changing proteins were selected by applying a log2-fold-change (>1.0) and an adjusted p-value (< 0.05) corrected for multiple testing threshold.

**Supplementary Figures:**

**Supplemental figure 1.** MYC Expression Correlates with Proscillaridin Anticancer Efficacy. **a** Upper panel, half maximal inhibitory concentration (IC_50_) after a 24h proscillaridin treatment (ranging from 1 nM to 100 µM) in a panel of human cancer cell lines (n=4). Lower panel, MYC protein level in each untreated cell line, assessed by western blotting. ACTIN was used as a loading control (n ≥ 3). **b** Graph showing MYC expression (relative to ACTIN) compared to proscillaridin IC_50_ (24h) in 14 cancer cell lines. Correlation was evaluated by linear regression analysis; P-value is shown on the graph (n=3). **c** Representative pictures of transformed primary human fibroblasts before and after transfection with *RAS^V12^*, *MYC* and *RAS*^V12^/*MYC* were taken by light microscopy (400X magnification). **d** Dose response curves and IC_50_ values after 48h proscillaridin treatment (ranging from 0.1 nM to 200 µM) in MOLT-4 (T-cell leukemia), NALM-6 (B-cell leukemia), and REH (non-T, non-B cell leukemia) (n=3). **e** Time course experiment where NALM-6 cells were treated with 5 nM for up to 96h. MYC expression was calculated as a ratio over ACTIN levels (*indicates P<0.05; ANOVA; n = 3).

**Supplemental figure 2.** Transcriptomic Analysis In MOLT-4 Cells Treated with Proscillaridin (5 nM, 48h). **a** Heat map representing RPKM similarities between triplicates of untreated (U) and proscillaridin-treated (5 nM; 48h; T) MOLT-4 cells (n = 3). Red color corresponds to the highest similarity and yellow corresponds to the lowest similarity. **b** Proscillaridin (5 nM, 48h) induced gene expression reprogramming of MOLT-4 cells. Volcano plots of gene expression changes in MOLT-4 cells in untreated versus treated samples. Black dots correspond to genes with P-value adjusted > 0.5. Grey dots correspond to genes with P-value adjusted < 0.5 but without significant fold change expression difference between untreated and treated cells (-0.5 < FC < 1). Downregulated genes with P-value adjusted < 0.5 and FC < -0.5 are shown in green. Upregulated genes with P-value adjusted < 0.5 and FC > 1 are shown in red. Numbers of downregulated and upregulated genes are shown on the graphs. **c** Metascape analysis of genes downregulated by proscillaridin treatment (5 nM; 48h). **d** Cell cycle analysis after BrdU staining in MOLT-4 and NALM-6 cell lines exposed to proscillaridin (5 nM, 48h). Cell fluorescence was measured by flow cytometry (* indicates P<0.05; ANOVA; n=3). **e** Metascape analysis of genes upregulated by proscillaridin treatment (5 nM; 48h).

**Supplemental figure 3.** Histone 3 Acetylation Loss Induced By Proscillaridin In MOLT-4 And NALM-6 Cells. **a** MOLT-4 cells were treated with proscillaridin (5 nM) and histones were acid-extracted after 8, 16, 24, 48, 72 and 96 hours. H3 acetylation levels were quantified and expressed as a percentage of untreated cells (* indicates P<0.05; ANOVA; n = 3). **b** Ratio of chromatin immunoprecipitation (ChIP) of H3K27 acetylation in MOLT-4 cells before and after proscillaridin treatment (5 nM; 48h) (*indicates P<0.001; t-test, n=3). **c** NALM-6 cells were treated with proscillaridin (5 nM) and histones were acid-extracted after 8, 16, 24, 48, 72 and 96 hours. H3 acetylation levels were quantified and expressed as a percentage of untreated cells (* indicates P<0.05; ANOVA; n = 3).

**Supplemental figure 4.** Histone 4 Acetylation Is Unchanged After Proscillaridin In MOLT-4 And NALM-6 Cells. **a** MOLT-4 and **b** NALM-6 cells were treated with proscillaridin (5 nM) and histones were acid-extracted after 8, 16, 24, 48, 72 and 96 hours. Histone 4 acetylation levels were assessed using antibodies against K5ac, K8ac, K16ac, K20ac, and total histone 4 acetylation. H4 was used as loading control. H4 acetylation levels were quantified and expressed as a percentage of untreated cells (* indicates P<0.05; ANOVA; n = 3). **c** Confocal microscopy (60X) of MOLT-4 cells stained with DAPI revealed heterochromatin modulation after proscillaridin treatment (5 nM; 48h). White arrows indicate loss of heterochromatin regions.

**Supplemental figure 5.** H3K27 Acetylation DNA Occupancy Is Lost After Proscillaridin Treatment In MOLT-4 Cells. Metascape analysis of **a** downregulated genes and **b** upregulated genes after proscillaridin treatment (5 nM; 48h) marked by H3K27ac in their promoter regions (-500 bp / +500 bp).

**Supplemental figure 6.** Proscillaridin Treatment Downregulated MYC Target Genes That Are Marked By H3K27ac In Promoter Regions. Map of **a** co-expression pathways and **b** protein-protein physical interactions of MYC target genes marked by H3K27ac in untreated MOLT-4 cells.

**Supplemental figure 7.** Loss of Acetylation In MYC Protein And MYC After Proscillaridin Treatment In High MYC Expressing Cells. **a** Mass spectrometry analysis on 2 MYC peptides (LVSEK(ac)LASYQAAR) after proscillaridin treatment (5nM; 48h) in MOLT-4. Log_2_ normalized intensity is shown (* indicates P<0.0001; t-test; n=4). **b** Map of co-expression pathways of the 28 proteins that lost acetylation after proscillaridin treatment (5 nM; 48h) in MOLT-4 cells.

**Supplemental figure 8.** MYC Inhibition Induced By Proscillaridin Is Regulated By KAT Activities. **a** MOLT-4 cells were treated with proscillaridin (5 nM) and histones were acid-extracted after 8, 16, 24, 48, 72 and 96 hours**.** KAT3A, KAT3B, KAT5, KAT2A, KAT2B, KAT6A and KAT7 expression levels were assessed by western blotting. ACTIN was used as loading control. **b** KAT2B and KAT7 expression levels were quantified and expressed as percentage of untreated cells (n=3). **c** MOLT-4 cells were treated with KAT3B/A inhibitor C646 (10 µM) and with proscillaridin (5 nM) and histones were acid-extracted after 48 hours. Histone 3 acetylation levels were assessed using antibodies against K14ac, K18ac, K27ac, and total histone 3 acetylation. H3 was used as loading control. **d** Left panel, MOLT-4 cells were treated with KAT3B/A inhibitor C646 (10 µM) and with proscillaridin (5 nM) and KAT5, KAT3A and KAT3B expression levels were assessed by western blotting. ACTIN was used as loading control. Right panel, KAT5, KAT3A and KAT3B levels were quantified and expressed as a percentage of untreated cells (* indicates P<0.05; t-test; n = 3).

**Supplemental figure 9.** Proscillaridin Induces KAT Downregulation Specifically In High MYC Expressing Cells. **a**, **b** and **c** MOLT-4, NALM-6, SW48 and A549 cell lines were treated with proscillaridin (5 nM, 48h) and fibroblasts transfected with *RAS^V12^*, *MYC* and *RAS^V12^/MYC* were treated with proscillaridin (5 nM or 70 nM, 48h). **a** KAT3A (CBP), KAT3B (P300), KAT5 (TIP60), KAT2A (GCN5), KAT2B (PCAF), KAT6A (MOZ) and KAT7 (HBO1) expression levels were assessed by western blotting. ACTIN was used as loading control. **b** Histone 3 acetylation levels were assessed by using antibodies against K9ac, K14ac, K18ac, K27ac, and total histone 3 acetylation. H3 total was used as loading control. **c** Histone 3 acetylation levels were quantified and expressed as percentage of control (* indicates P<0.05; t-test; n=3).
